# Perceptual constraints of motion shape animal movement strategies

**DOI:** 10.64898/2026.09.22.753148

**Authors:** Violette Chiara, Samuel R. Matchette, Pritish Chakravarty, Nadia M. Hamilton, Palina Bartashevich, Joanna R. Attwell, Christos C. Ioannou, James E. Herbert-Read

## Abstract

As animals explore their environment, self-generated motion can affect visual perception, while perceptual constraints shape movement decisions. Movement strategies should therefore balance the benefits of exploration against the potential perceptual costs of motion. Here, we test whether the movement strategies animals adopt can enhance information acquisition while accounting for the constraints that motion imposes on visual perception. We first quantified the visual perceptual abilities of freely moving fish in three dimensions by projecting virtual prey around three-spined sticklebacks (*Gasterosteus aculeatus*), determining how the likelihood of detecting prey varied across the fish’s visual field as a function of their movement. Self-induced motion reduced the likelihood prey were detected, with increases in speed non-linearly reducing the perceptual abilities of the fish. Using agent-based simulations based on empirical models of visual perception, we demonstrate that when visual perception degrades non-linearly with speed, the strategy that maximizes prey detection is ‘saltatory’ locomotion, involving stationary pauses interspersed with movements. Moreover, experiments with freely swimming fish showed they adopted specific durations of stops and movements that improved prey detection given the way motion impacts their perception. Our results reveal that some movement strategies are more effective than others at gathering information during exploration.

## Introduction

The ability of an animal to detect resources and risks in its environment often determines the outcome of inter- and intra-specific interactions, ultimately impacting ecological and evolutionary processes such as predator avoidance, mate selection or foraging success [1]. For many animals, the information they detect while exploring shapes their subsequent movement decisions [2]. However, an animal’s movement can also impact its ability to detect information. For instance, self-induced movements can cause motion blur, where objects appear blurred in the visual field owing to long response times of photoreceptors [3, 4]. Despite eye and head movements compensating for some degree of motion blur [5, 6], self-induced motion can affect an animal’s ability to detect small or distant objects against a background [7-10], which may be particularly important for tasks such as detecting prey. This phenomenon of movement-induced sensory impairment is not limited to the visual domain: olfaction [11], mechanoreception [12], and the lateral line [13] can also be negatively impacted by self-induced motion.

If perception is compromised on the move, then animals are predicted to adopt movement strategies that mitigate these effects [9, 14-16]. Moreover, which movement strategies animals adopt are predicted to be determined by how movement impacts perception. Anderson [9] predicted that animals should use a ‘cruising’ strategy, with animals moving at a constant speed without stopping, if increases in speed led to proportional decreases in perception. Alternatively, if motion impacts perception non-linearly, with small increases in speed having a disproportionally large reduction in perception, then saltatory motion, in which an animal moves in bouts that are interspersed by stopping phases, is predicted to be the most likely movement strategy to improve information gathering while exploring [9, 14, 16]. Indeed, this form of start-stop motion is commonly observed in many taxa, from unicellular organisms through to birds and mammals [17]. However, our understanding of the functional benefits of saltatory motion is still dominated by theoretical models, which generally rely on assumptions about animal perception rather than direct empirical measurements (e.g., [4, 9, 18, 19]). Indeed, while the stopping phases of saltatory motion are predicted to alleviate the perceptual cost associated with movement [9], whether the perceptual effects of self-induced motion link to movement strategies remain to be empirically tested.

Traditional approaches to estimating the theoretical limits of an animal’s visual perception use the anatomical position of the eyes, or histological measurements of the spatial arrangement of photoreceptors on the retina [20-24]. But to estimate the visual perception of animals on the move, specific behavioural responses to objects in the visual field can be used as a proxy for perception [25-27]. Here, we use the detection of virtual prey by freely moving three-spined sticklebacks (*Gasterosteus aculeatus*) in a 3D augmented reality environment to quantify how stickleback visual performance degrades as a function of self-induced motion. We then incorporate an empirically derived model of the visual perception of these fish into an agent-based simulation and compare prey detection rates of simulated fishes with different movement strategies. Based on these empirical data and theoretical investigations, we test whether the movement strategies adopted by individuals maximizes their ability to detect prey given the constraints that self-induced motion places on perception.

## Materials and Methods

### Prey detection experiment

To estimate the visual perception of freely swimming fish (sticklebacks, see Section 1.1 of Supplementary material), we used an augmented reality environment to project virtual prey while tracking the movements and responses of the fish in 3D (Fig. 1). The experimental arena consisted of a glass tank (55 x 40 x 35 cm; L x W x H). All internal glass walls except one were covered by opaque white plastic (Correx). On the internal wall that was not covered by Correx, we inserted a frame with a translucent screen attached (Rosco gel No. 252) so that it contacted the glass wall. Virtual prey (red dots, 3 mm in diameter) were projected through the glass and onto this screen using a projector (Benq MW523), positioned 82 cm outside the tank. The translucency of the screen allowed prey to be visible to the fish on the internal wall of the tank, while also allowing the fish to be visible through the screen. Above the arena, we angled a mirror at ∼45°. We positioned a camera (Panasonic 4K HC-VX870) facing the screen so that this wall of the tank and the mirror were in its field of view. This allowed us to capture two planes of projection through a single camera. The two planes were approximately orthogonal to one another (i.e., a horizontal plane XY, Fig. 1A, and a vertical plane XZ, Fig. 1B). Trials were filmed at 3840 x 2160 resolution and 25 frames per second. The camera was controlled remotely using the Panasonic Remote Control application. The arena was lit by fluorescent lamps placed vertically at two ends of the tank.

**Figure 1.**
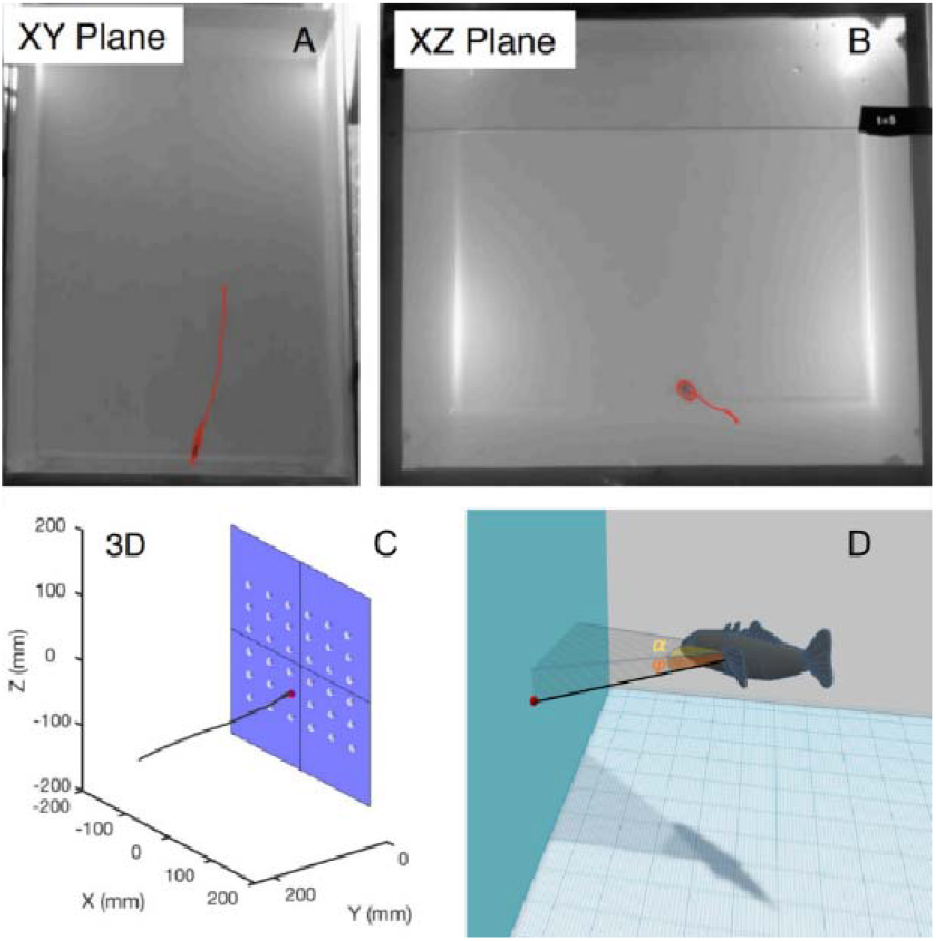
The augmented reality environment and methodology used to infer the visual perception of freely moving fish. A single frame from a video showing the (A) horizontal (XY) and (B) vertical (XZ) plane of the arena, which were captured simultaneously. Tracks in red represent positional and orientational tracks of the fish in both planes. (C) 3D reconstruction of the trajectory of the fish. The white-dotted grid represents the 36 positions where the prey could appear on the vertical side of the arena. (D) Illustrations of the azimuth (*α*) and elevation (*φ*) angles between the fish’s orientation and the position of the virtual prey. The red dots represent the position of the virtual prey.

We used an open-loop feedback system built in MATLAB (version 2018a) to project single prey items in one of 36 positions on to the translucent screen (arranged in a gridded array; see Fig. 1C). To do this, we used a webcam (Logitech C615 HD) to view the vertical plane (XZ) of the tank on a computer monitor in real-time. Using a bespoke algorithm written in MATLAB, the experimenter randomly selected a point on the monitor, and the algorithm would determine the closest of the 36 projection points to the user-selected point. A single virtual prey then appeared at that location and remained present for 0.88 seconds, after which it disappeared. We chose a relatively short presentation time to reduce the likelihood of the fish learning the prey were not edible. Because prey motion typically facilitates detection by predators, the prey was programmed to move at a speed of ∼1 cm/s on a correlated random walk, but in total moved < 1 cm from its starting position before disappearing. The starting position that each prey appeared at was saved. At the time the prey was projected, a counter (only visible from outside the arena) was projected onto the external wall of the arena that showed the prey number. This counter allowed us to subsequently identify when each prey presentation occurred, and later the fish reaction to it.

For each trial (n = 100), an individual stickleback was placed in the arena and left to acclimate for 10 minutes. After these 10 minutes, we began filming and projecting the virtual prey. Each stickleback received 50 prey presentations over the course of its trial. Trials usually lasted ∼10 minutes from the first prey presentation until the last, with the specific trial length dependent on when the prey were projected. The minimum time between consecutive prey presentations was at least two seconds. Individual fish were never used more than once.

After each day of trials, we performed a camera calibration procedure so that we could determine the real-world coordinates of both the virtual prey and the fish. To do this, we first projected the 36 possible locations of the prey as white dots onto the screen, allowing us to record their pixel coordinates in both planes. We also placed a calibration board (7 x 9 black-and-white checkerboard with 30 mm x 30 mm squares) underwater in the arena so that it was visible in both the filming planes (camera and mirror: Fig 1A, B). We captured 30 images of this calibration board in different orientations and positions in the arena. Using these images, we used MATLAB’s Camera Calibration toolbox to identify the positions of the camera and mirrored camera view in the scene, and to construct camera calibration matrices for conversion of measurements into real-world coordinates.

We tracked the mid part of the body and the orientation of the fish from one second before to two seconds after each prey presentation using CTrax [28]. We subsequently used the FixErrors GUI in MATLAB to correct errors produced by the tracking software. We used the camera calibration matrices to convert these trajectories and the possible locations of the prey into the real-world 3D coordinates (Fig. 1C). Recordings from the lateral side (vertical plane XZ) were also watched by an experimenter who identified for each prey presentation whether the fish reacted to the prey or not (see [29, 30]). For each prey presentation, we manually scored whether the fish reacted to the prey (red dots are perceived as food; see [29, 30]) by identifying stereotypical changes in behaviour, including an acceleration or change in direction of the fish towards the prey ([31], see Fig. S1). We considered that a reaction occurred if the fish changed its orientation and redirected its movements toward the prey before the prey disappeared (Fig. S1 shows quantitative differences in the behaviour of fish that reacted to the prey or not). Because the frame used to attach the translucent screen sometimes obstructed the view of the fish on the video at the edges of the arena in the vertical plane (XZ), making tracking and fish reaction difficult to estimate, we removed some prey presentations from the analysis. In total, we corrected the trajectories of the fish in 4637 prey presentations (out of a possible 5000) in both planes.

Using the fish trajectories and prey positions, we first asked how the likelihood of responding to the prey was affected by where the prey appeared in the fish’s visual field and the speed of the fish. Using bespoke scripts in MATLAB, we determined where the prey appeared in the fish’s visual field in the X, Y and Z axis, relative to the fish located at the origin (X = 0, Y = 0, Z = 0) and facing along the positive X axis. These data were then imported into R (version 4.2.1) to extract different variables measured at the frame the prey appeared. We first calculated the speed of the fish as the distance travelled between consecutive frames, converted to mm/s. We estimated the azimuth angle between the fish and the prey (*α*), which is the angle between the vector defining the fish’s heading and the vector between the fish’s and the prey’s location in the horizontal plane. This value varied between -180° (behind and to the left) and 180° (behind and to the right) (Fig. 1D). We also calculated the elevation angle (*φ*) between the fish and the prey. This angle reflects the angle between two vectors, one that runs from the body of the fish to the arena wall matching the pitch of the fish and one that runs from the mid-body of the fish directly to the location of the prey (Fig. 1D). The angle (*φ*) ranges from -90° (prey directly below the fish) to 90° (prey directly above the fish). For later analyses, we used the values of these variables corresponding to the first frame for which the prey appeared.

### Statistics

To model the likelihood that fish would detect prey as a function of where the prey appeared in their visual field and how fish were moving, we used a binomial Generalized Linear Model (GLM), with the reaction to the prey (1 if there was a reaction, 0 otherwise) as the response variable. Fixed effects included: i) the distance between the midpoint of the body of the fish and the prey when projected on a horizontal plane (XY), ii) either a second-order polynomial (poly from the *stats* package) of the azimuth angle (*α*) between the fish and the prey or the absolute value of this angle, iii) a second-order polynomial (poly from *stats* package) of the elevation angle (*φ*) between the fish and the prey or its absolute value, and iv) the speed of the fish (Fig. 1D). We did not include the individual’s identity as a random factor in the model, because the resulting models were too complex and led to convergence problems. Given the substantial number of tested individuals (n = 100), with each individual undergoing an equal number of prey presentations, individual variation is unlikely to have contributed substantially to model outputs.

All variables were standardized (subtracting the mean and dividing by the standard deviation), and models were compared using Akaike’s Information Criterion (AIC). The most complex models included all possible two-way interactions. We then used the model with the smallest AIC (see supplementary Dataset S1) to predict the probability of prey detection in various conditions using the predict function from the R package *stats*. The assumptions of the models were verified using the R package *DHARMa* [32]. Models were checked for overdispersion with the testDispersion function (GLM: P = 0.78) and associated Kolmogorov–Smirnov tests (simulateResiduals function) (GLM: P = 0.26, GLMM: P = 0.34). To check for outliers, we used the function testOutliers with type = “bootstrap” because of the binomial nature of our data (GLM: P = 0.5, GLMM: P = 0.38). All statistical analyses were carried out in R (version 4.2.1), and data are available in Dataset S1. This model was used as a proxy for the fish’s visual perception. We checked the overall importance of the final terms included in this model through additional model comparison approaches [33] (see *SI Appendix* section S1.2, Equation S1, and Fig. S2, S3).

### Agent-based model simulations

To find the movement strategies that maximized the number of detected prey given a specific perceptual model, and compare model predictions with experimental data, we developed a three-dimensional Monte-Carlo agent-based model in Python (Python 3.9). The simulation is parameterized using a physical metric scale, such that coordinate values correspond directly to real-world distances. Distances and velocities are therefore reported in standard metric units (e.g., mm, m, mm/s). This model simulated a fish moving along a straight line in a 3D environment for a duration of three minutes. We used a time step of Δ_t_ = 1/23.98 sec similar to the frame rate used for the previously described experimental recordings. Prey were presented in 3D around this line with a prey density set at 0.5 prey per m^3^. Prey positions were randomly assigned around the fish’s path (maximum distance of prey to the path = 2 m). Prey were motionless and present from the beginning of the simulation. They would disappear only when detected by the fish. The total length of the line was calculated so that the fish would never reach its end, even if moving constantly at maximum speed (total length was the length travelled by the fish at maximum speed during 3 minutes plus 4 meters). Therefore, the length of the path was variable between the simulations, adapted to the fish movement parameters, and the number of prey was adapted accordingly to ensure the same prey density between simulations.

The simulations involved four parameters, which were informed by experimental parameters from the exploration assay (see below): mean duration of motion bouts (movement episodes consisting of acceleration, constant non-zero speed, and deceleration phases), mean duration of stops (fish moving speed equals zero), speed during motion bouts, and the shape of the perceptual decrease with increasing speed. This last parameter had three options (Fig. S4): convex decrease (based on our perception model from the experimental data), linear decrease (identical to the first but corrected by a linear/convex ratio so that the overall shape of the decrease follows a straight line), and motion blindness (there was zero probability of the fish detecting prey during motion, and the same probabilities as the ones calculated from empirical data when motionless). When prey was located less than 2 m from the fish, we determined whether it was detected or not based on the probabilistic perception model (see *SI Appendix*, section S3.1). Prey remained present until the end of the simulation or in case of detection by the agent fish. If prey were never removed from the simulations, fish could stay motionless and repetitively detect the same prey, something that would make little biological sense. Therefore, prey that were detected were immediately removed from the simulation and did not reappear, so as not to be detected twice. For simplicity, the agent’s behaviour was not altered upon detection of, or failure to detect, prey. Several prey can be detected in one time step.

To quantify prey detection across different movement strategies and perceptual degradation models, we conducted simulations in which we systematically varied four key parameters: average duration of movement phases (0, 0.5, 1, 2.5, 5, 7.5, 10 s), average duration of stopping phases (same values), movement speed (50–400 mm/s in 50 mm/s increments), and perceptual shape function (convex, linear, motion blindness). For each unique combination of parameter values, we ran 5,000 replicate simulations, resulting in a granular grid of scenarios (see Fig. S5). For each simulation, the total number of prey detected was calculated, and this value was then averaged across the 5,000 replicates (see SI Appendix, Section S3). This initial simulation set provided an approximation of how detected prey number varied across the multidimensional parameter space. To more precisely estimate the parameters that maximized prey detection for a given perceptual model, we employed an iterative grid refinement procedure [34]. First, we identified the motion speed associated with the maximum average prey detection rate in the coarse grid – we use the term ‘optimal’ in the sense of maximizing prey detection rate. We then constructed a finer-resolution parameter grid centred on the optimal values found, setting speed constant at the optimal speed found in the preceding step, and varying the duration of motion bouts and stopping phases at finer resolution. Because the number of prey detected was highly variable between simulations, and because of the very fine scaling of parameters, we averaged estimates over 10,000 replicates for the same parameter combinations. This process was repeated iteratively, refining grid resolution at each step until the resolution of the grid reached 0.125 s (∼ 5 to 10 times smaller than the average moves duration observed in empirical data) for motion bouts and stopping phases durations. This allowed us to identify optimal movement strategies for prey detection with high precision (see Fig. S6).

### Exploration assay

To test whether real fish adopted movement strategies that maximized their theoretical likelihood of detecting prey given their empirically-derived perceptual model, we used a new batch of wild sticklebacks, separate from those in the virtual prey experiment (see ESM, Section 2.1). Trials were conducted between June and October 2021. Each trial (n = 162, 81 fish, with each fish tested twice) comprised of an individual fish being placed in a circular arena (80 x 29 cm; diameter x height) with a depth of 15 cm. Positioned above the arena was a camera (Nikon D7000; 1920 X 1080 pixels @ 24 fps) to film each trial, and a projector (ELEPHAS Projector Q9 Native 1080P HD) to project the stimuli backgrounds. Backgrounds were chosen to appear more naturalistic and therefore promote explorative behaviour [35]. Black curtains were fitted around the experimental set-up to minimize visual disturbances. At the start of each trial, an individual was transferred from its individual mesh container to the experimental arena for a 10-minute acclimation period. During this acclimation phase, individuals were exposed to a greyscale background projection (RGB: 195, 195, 195), mimicking the average RGB values of the experimental backgrounds. Following the acclimation phase, the first experimental background stimulus was slowly faded in (for a duration of 10 seconds) to avoid startling the fish. This background remained for 10 minutes, at which point a second transition occurred, whereby the second experimental background faded in over the first. Within each trial, the analysis of movement parameters excluded the acclimation and the two transition periods. Following a trial, the focal individual was returned to its individual mesh container, and the process repeated for the next individual. Both stimuli backgrounds comprised of the same image of a rocky substrate, with one at 100% magnification and the other at 200% magnification (Fig. S7). The order in which each fish received each background was alternated across trials. All individual fish completed two trials, with a 4-day interval between the two.

We used the software Loopy (LoopBio) to extract the XY trajectories for each individual fish from the videos. This was achieved using a supervised machine-learning approach, whereby manually annotated frames were used to train a “key point detector” model to identify the head and tail of each fish in a given video. For training, a total of 960 frames from 20 videos (48 frames per video) were manually annotated. Once trained, all videos were passed through the model, and the subsequent XY movement tracks were extracted. The tracking results were then imported into AnimalTA [36] for manual correction. All videos were corrected by the same experimenter.

From these tracking data, we aimed to identify four different movement phases: fish was moving, fish was stationary, fish was accelerating, fish was decelerating (see Fig. S8). These phases were identified using a bespoke R script (v 4.2.1) [37]. It was sometimes impossible to correct missing coordinates as the fish were hidden by a reflection or were otherwise not visible for a short period of time. In that case, only uninterrupted trajectories of at least 5 seconds were kept for later analyses. To identify acceleration and deceleration phases, trajectories were smoothed using Savitzky-Golay filtering with a window length of 5 frames and a polyorder of 1 (function sgolayfilt from the signal package). The acceleration or deceleration value (i.e., the difference in speed between two frames) was then calculated and these values were smoothed with the Savitzky-Golay filtering (window length of 3 frames, polyorder of 1). Accelerations and decelerations were defined as periods where the change in speed exceeded 1 mm/s per frame and that lasted longer than 1 frame (approximately 0.042 seconds). In addition, these measurements were only considered if the minimum speed was lower than 15 mm/s and the maximum speed was higher than 15 mm/s. These thresholds were selected based on visual inspection of the data. Movement phases were identified as phases between acceleration and subsequent deceleration periods, while stopping phases were phases between deceleration and subsequent acceleration periods (see Fig. S8). Phases between two consecutive accelerations or two consecutive decelerations were uncommon and were excluded from the analysis. We used paired Wilcoxon-rank tests to assess whether there were major differences in these behaviours between the two treatment backgrounds; the duration of stops (V = 1931, p = 0.20) and movements (V = 1866, p = 0.33) did not differ between backgrounds, nor did the speed while moving (V = 1646, p = 0.95), therefore we analysed average movement parameters across the two different backgrounds together. The patterns of acceleration and deceleration were used to inform the acceleration and deceleration profiles in the agent-based simulations, while the duration of moving and stopping phases, along with the maximum speed during a movement phase (before smoothing), were used to compare between experimental and simulation data.

## Results

### Visual perception model

Fish in the augmented reality environment responded to 7.18% of the 4637 usable virtual prey presentations, with a range of 0-16 reactions per fish. Analysis of fish and prey coordinates, together with fish behavioral responses, indicated that fish were more likely to detect prey that appeared directly in front of them (azimuth, *α* ≈ 0 degrees), with the likelihood of detecting the prey decreasing if prey appeared in lateral regions of the visual field (i.e. *α* ≫ 0 degrees) or when prey were further away (Fig. 2A and Movie S1). The best fitting model included an interaction term between the distance to the prey and the azimuth angle, *α*, at which the prey appeared in the visual field (LRT = 11.14, df = 1, p < 0.001; Table S1; Fig. 2A). If prey appeared directly in front of a fish (*α* ≈ 0 degrees), the decay in detection probability with distance was reduced compared to prey that appeared in lateral regions of the visual field (*α* ≫ 0 degrees), where the decay was more rapid (Fig. 2A).

**Figure 2.**
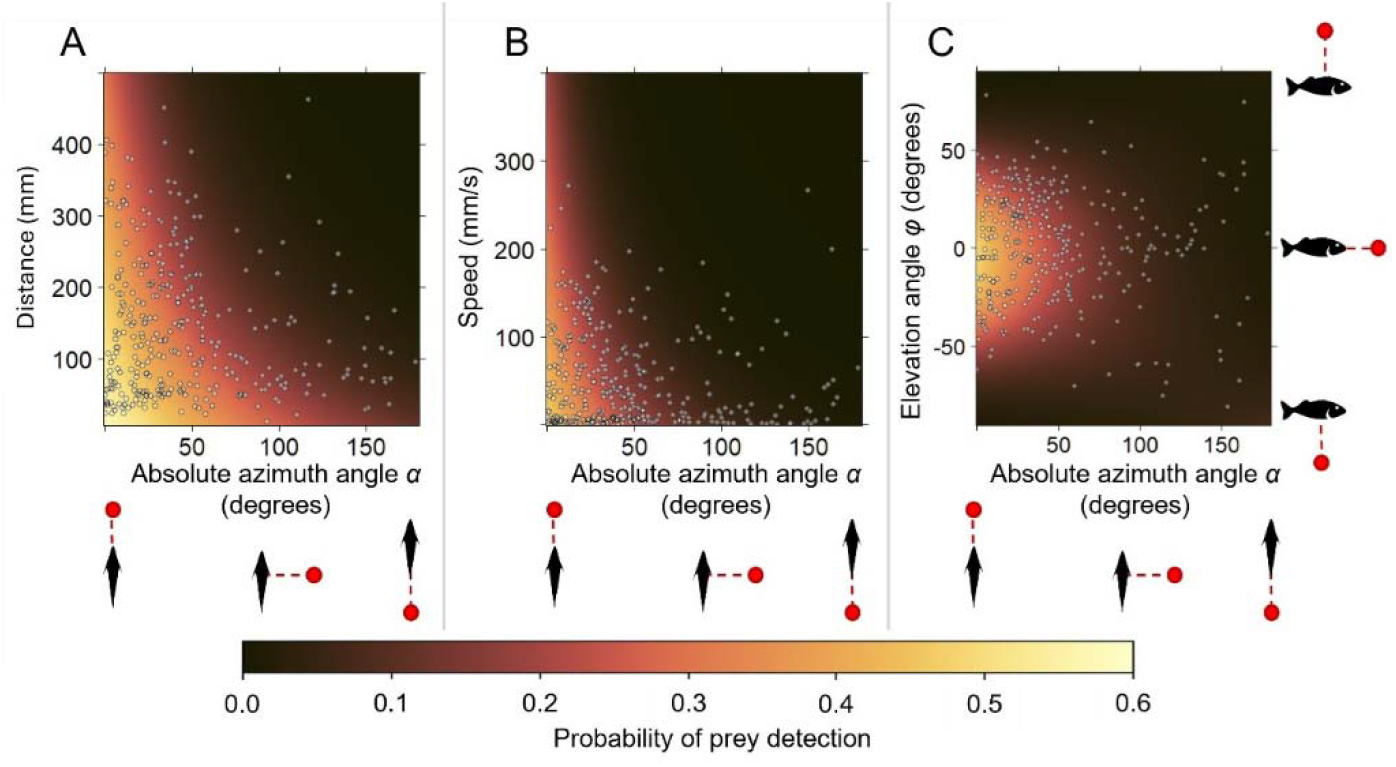
Heatmaps illustrating significant interactions in the GLM used to model the visual perception of fish. (A) Estimated probability of the fish detecting the prey as a function of distance to prey and absolute azimuth angle, α. For these predictions, the elevation angle, φ, was set to 0° and speed was set to 0 mm/s. (B) Estimated probability of the fish detecting the prey as a function of the fish’s speed and absolute azimuth angle, α. For these predictions, the elevation angle φ was set to 0° and the distance to prey taken as the average observed (180.9 mm). (C) Estimated probability of the fish detecting the prey as a function of the fish’s elevation angle φ and absolute azimuth angle, α. For these predictions, speed was set to 0 mm/s and distance to prey taken as the average observed (180.9 mm). Points represent experimental events (n = 333) where fish detected the prey. Events without detection are not represented for clarity. Both distance to prey and the speed of the fish reduced the likelihood of fish detecting the prey and reduced the visual field (azimuth angle α) over which prey were detected.

The speed that the fish were moving at impacted the regions of the visual field where fish detected the prey (interaction between speed and azimuth *α*; LRT = 12.16, df = 1, p<0.001; Table S1; Fig. 2B). With a speed change from 50 to 150 mm/s, fish were 17.8% less likely to detect prey when the prey were directly in front of them (prey at *α* = 0°, φ = 0°, 15 cm distance), whereas they were 55.4% less likely to detect prey that appeared more to their side (prey at *α* = 45°, φ = 0°, 15 cm distance). In effect, travelling at increased speed induced a “tunnelling effect” [38], where the region directly in front of the fish remained the primary region where prey could be reliably detected. The visual perceptual model also included an interaction term between the second order elevation angle φ and azimuth angle *α* (interaction term between second order elevation φ and azimuth *α*: LRT = 10.28, df = 2, p = 0.006; Table S1). This interaction term indicated that changes to the elevation angle had larger impacts on prey detection for smaller rather than larger azimuth angles. For example, when prey were in front of the fish (prey at *α* = 0°, speed = 0 mm/s, 15 cm distance), changes to the elevation angle from 0 to 50 degrees (prey being above the fish head) reduced prey detection by 70.7% compared to only 57.6% when prey appeared to the side of the fish (prey at *α* = 90°, speed = 0 mm/s, 15 cm distance) (Fig. 2C). Prey detection also differed according to elevation, with prey presented at positive elevation angles (prey above the fish’s head) being on average 30.6% less likely to be detected than prey presented at negative elevation angles (average probability of detection for prey at 15 cm distance and moving speed of 0 mm/s).

This statistical perceptual model (see *SI Appendix*, section S1.4 and Equation S2) allowed us to produce three-dimensional volumetric regions which depicted the likelihood that fish would detect prey as a function of where they appeared in the visual field and the speed the fish was moving. These regions highlighted that the perception volume of the fish is more complex than a simple sphere or ellipsoid, and is altered by the fish’s speed (Fig. 3A-B). Increases in speed reduced the volume over which prey were detected with a given probability (Fig. 3A-B). The overall decrease in perception with speed was characterized by a convex curve (Fig. S4), with the likelihood of detecting prey (see *SI Appendix*, section S1.2) decreasing sharply with increasing speed before plateauing.

**Figure 3.**
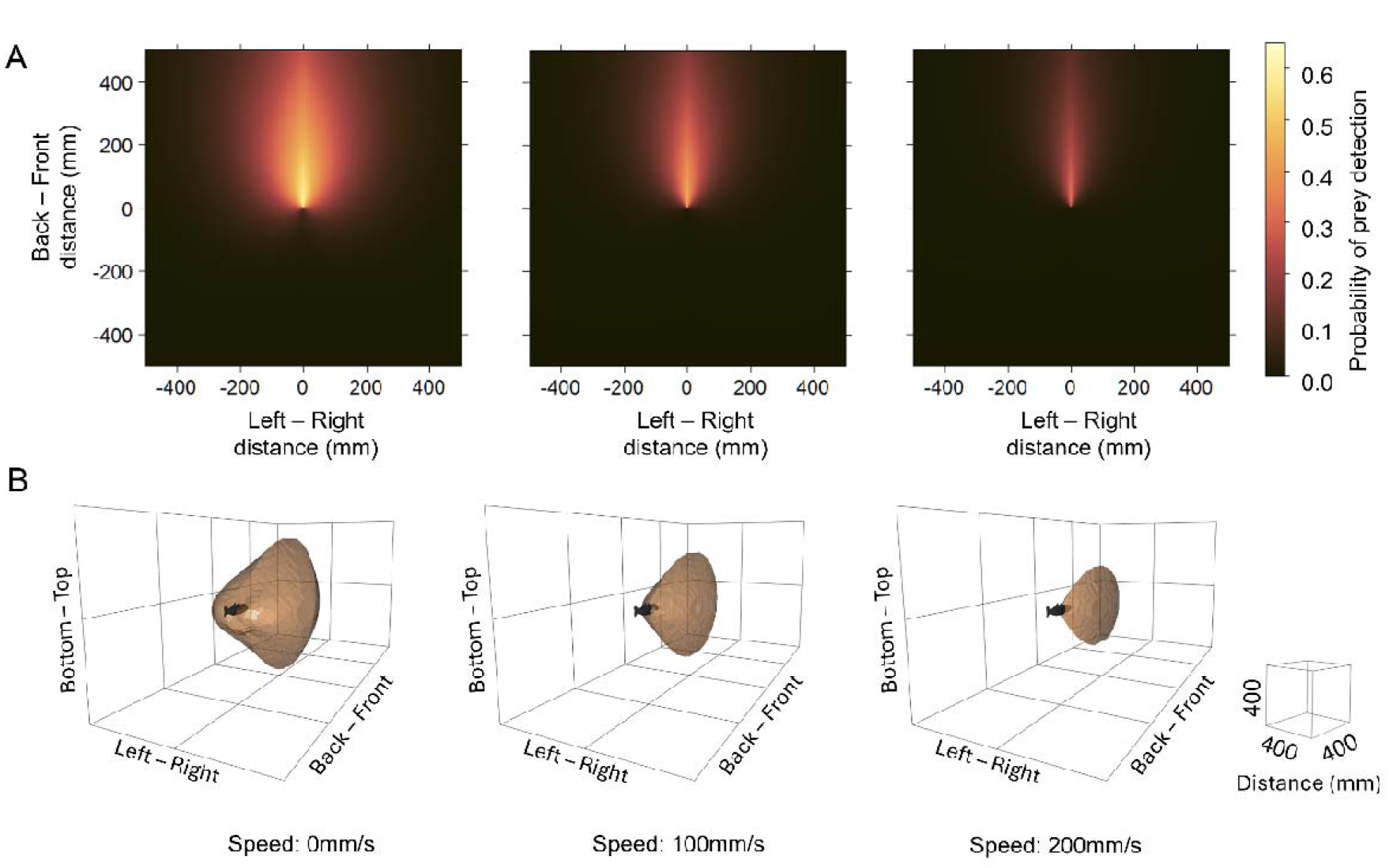
Illustration of how speed impacts the perceptual ranges of fish. (A) The probability of fish detecting the prey (predicted using the GLM model) according to the relative position of prey with respect to a fish with mid-body located at (0,0) and facing along the positive Y axis.

Probability maps are shown for three movement speeds (left: 0 mm/s; centre: 100 mm/s; and right: 200 mm/s). Maps were created for prey on the horizontal plane of the fish (Z = 0). (B) Representation of the regions around a fish where the probability to detect a prey within the shaded volume is 0.15 or higher. The shape and volume *V* of this field vary according to the fish’s speed *s* (left: *s =* 0 mm/s, *V* ≈ 94.46 dm^3^; centre: *s =* 100 mm/s, *V* ≈ 34.33 dm^3^; and right: *s =* 200 mm/s, *V* ≈ 11.17 dm^3^), the volumes have a vertical symmetry plan (left and right parts relative to fish orientation are identical), but no horizontal plan of symmetry (volume above and below the fish are different). The schematic indicates the fish’s position and orientation, although for illustrative purposes, the fish is shown at four times its actual size.

### Movement strategies as a function of perceptual constraints

Anderson [9] predicted that saltatory motion should be used to improve information detection on the move if individuals have a non-linear decrease in their perceptual abilities as a function of speed. We tested Anderson’s predictions, asking whether fish with our empirically-derived perceptual model improved information detection with saltatory motion compared to other movement strategies. Our three-dimensional, spatially explicit agent-based model simulating a fish moving was first used to model an agent having a perceptual model as derived from the empirical data described above, with the overall probability of prey detection decreasing with speed following a convex curve (red line in Fig. S4; Fig. S3). With this convex perceptual model, the movement strategy that maximized prey detection was saltatory motion, with agents moving with short stops and short moves (Fig. 4A, Fig. S5), supporting Anderson’s predictions. With this convex perceptual model, faster speed during motion bouts increased the number of prey detected (Fig. 4A, Fig. S5). When the perception of agents was modelled to degrade linearly with increasing speed (blue line in Fig. S4; Fig. S3), the simulations instead predicted cruise search as the best movement strategy, with agents moving with constant speed without stopping (Fig. 4A, Fig. S5). The form of the linear decrease in perception with speed could be modelled with various slopes and intercepts, but cruise search remained the most advantageous strategy regardless of these modelling choices (Fig. S9). If, however, agents were modelled to be effectively blind while moving, as has been suggested to occur in some animals [39, 40], saltatory motion was the most advantageous strategy. In particular, when the likelihood of detecting prey fell to zero irrespective of speed attained during motion bouts (green line in Fig. S4; Fig. S3), a combination of stops and movements maximized prey detection rates.

**Figure 4.**
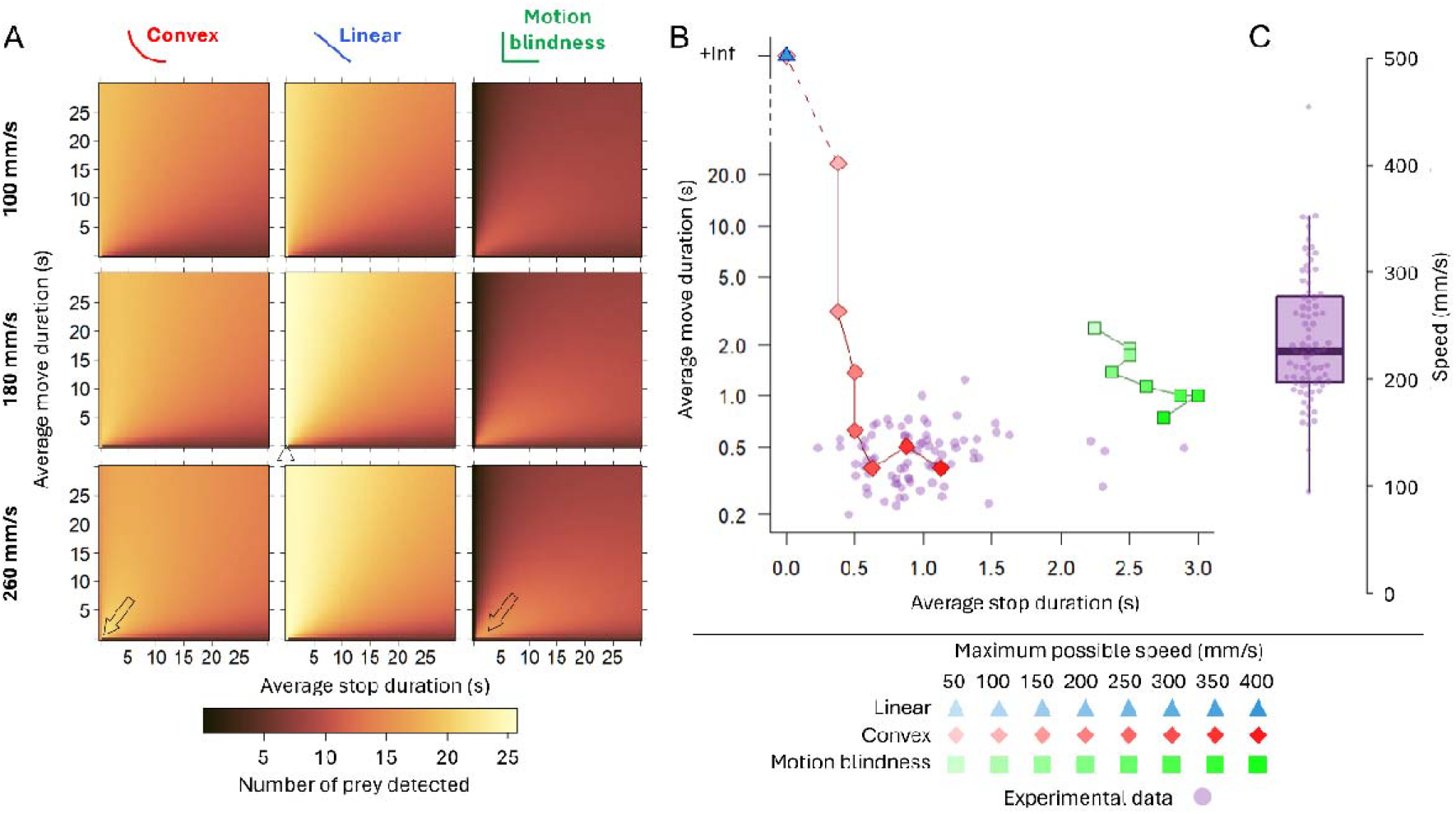
Simulated and empirical moves–stop dynamics and their effects on prey detection across perceptual models (A) Number of prey detected in simulations of moving agents (n = 5000 per grid cell). The resolution of each grid is 1 cell per 1 second for both moves and stops durations. Locomotion ranged from purely stationary (duration of movement = 0 s) to purely “cruising” (duration of stops = 0 s). Except during acceleration and deceleration, the agent’s speed while moving was fixed to 100, 180 or 260 mm/s. Each column of plots in the subfigure represents one of the three perceptual models. In the left column, the decrease in perception with increased speed follows a convex curve, as observed experimentally. In the central column, the agent’s perception decreases linearly with speed. The right column describes agents that cannot detect prey while moving. The arrow in each column indicate the location within parameter space that resulted in the maximum number of detected prey with the associated perceptual function. In the case of the linear perceptual decrease, the arrow indicates that the strategy with the maximum number of prey detected prey occurs when the average stop duration = 0, i.e. a cruising strategy. (B) Average durations of motion bouts and stopping phases extracted from experimental data (purple, n = 81) and combinations of these parameters predicted by our models (linear perception model: blue triangles, convex: red diamonds, motion blindness: green squares) that maximize prey detection rates (see *SI Appendix*, sections S2 and S3; Fig. S3 and S9) for various maximum speed limits (indicated by color intensity). When the average stop duration = 0 s, the fish continually moves (Y-axis) and adopts cruise search. Note that for the linear perception model, the movement strategy that maximizes the number of detected prey is cruise search. For this reason, the eight points are overlaid, resulting in only one visible point. Durations of motion bouts and stopping phases predicted to maximize prey detection rates based on the convex perception model best match those adopted by the real fish (purple points). If fish are blind while moving, stop durations are expected to be longer than those observed empirically. A logarithmic scale is used for the Y-axis to improve readability. (C) Average movement speed of the fish measured experimentally (see *SI Appendix*, section S2). The box represents the interquartile range (IQR), the line inside the box indicates the median, and the whiskers extend to the minimum and maximum values within 1.5 IQR.

While the convex and motion-blindness perceptual models both predicted saltatory motion as the strategy that maximized prey detection, the motion parameters that maximized prey detection rates differed between the two models. In particular, in the motion-blindness perceptual model, the most advantageous strategy was for agents to stop for longer periods of ∼ 2.9 s before moving at a maximum speed for ∼1 s (Fig. 4A-B, Fig. S5-6). On the other hand, in the convex perceptual model, the duration of stops and motion bouts that maximized prey detection was shorter, at 0.63 s and 0.38 s at maximum speeds, respectively.

### Comparison between model prediction and exploration assay

Freely swimming sticklebacks tested in the exploration assay adopted a saltatory movement strategy, with stops interspersed with motion bouts at speed (Fig. S8). Moreover, the average stop and move duration of those real fish (Fig. 4B) aligned with the movement parameters predicted to maximize prey detection given the convex perceptual model. The duration of stops and movements that maximized prey detection rates with saltatory motion decreased as the maximum speed of agents in the model was allowed to increase (Fig. 4B). At speeds that real fish tended to adopt (mean maximum speed during motion bouts was 226 ± 34 mm/s; Fig. 4C), there was strong agreement between the movement parameters of the models that maximized prey detection rates and those adopted by real fish (Fig. 4B). In particular, real fish had stops that lasted for 0.99 ± 0.29 s, and motion bouts that lasted for 0.47 ± 0.16 s. Together, these findings support theoretical predictions that fish will adopt movement strategies that maximize their prey detection rates given the constraints that self-induced motion places on their perception.

## Discussion

We provide empirical evidence that the movement strategies animals adopt can be shaped by the sensory constraints imposed by self-motion, particularly in the visual domain. By integrating three-dimensional behavioural assays with virtual prey presentations and agent-based simulations, we demonstrate that the saltatory movement strategy observed in three-spined sticklebacks (*Gasterosteus aculeatus*) likely serves to maximize information gathering, specifically under the constraint that visual perception degrades non-linearly with speed.

Previous studies have suggested the existence of a relationship between movement strategy and perception [9, 18], but direct empirical tests have been lacking. Our findings show that small increases in swimming speed can impair visual sensitivity in sticklebacks, especially in peripheral regions of the visual field. This is partly consistent with the phenomenon of motion blur [4, 5], where higher speeds result in a form of sensory ‘tunnelling vision’ [40]. However, the degradation of perception followed a convex function, where the cost of movement on perception degrades sharply with increasing speed. This does not fully capture the effects of motion blur at lower velocities, where blur is typically negligible because photoreceptors can respond rapidly relative to the slow movement of images across the retina. As speed increases, however, images drift across the retina faster than photoreceptors can effectively track, while saccadic eye movements become less capable of compensating for this displacement. Consequently, motion blur becomes increasingly pronounced and may begin to impair visual performance as speed increases [5]. If motion blur were the sole perceptual constraint associated with movement speed, we would therefore expect a concave decline in perceptual performance with speed. Instead, the convex pattern observed in our results likely reflects interactions between motion blur and additional mechanisms affecting perception at higher speeds. For example, moving animals tend to orient their gaze and visual attention toward the region ahead of their trajectory, enabling anticipation of future paths and avoidance of obstacles [41]. In humans, studies further suggest that cognitive processes may narrow attentional focus toward events occurring directly ahead during locomotion [38, 42]. Such mechanisms would be expected to produce a rapid reduction in peripheral visual processing with increasing speed, consistent with our findings. An additional explanation for the general decrease in detection capacity is cognitive–motor interference, where humans face a trade-off between movement speed and cognitive task performance [43, 44]. To our knowledge, however, these phenomena have not yet been described in non-human species. Regardless of the underlying mechanisms of the observed perceptual decrease, this non-linear perception decrease supports theoretical predictions by Anderson [9] that saltatory motion is the optimal strategy when perception is degraded non-linearly with speed.

Saltatory motion is widespread across taxa, yet explanations for this behaviour have often focused on the energetic impacts [14, 45, 46] or risk reduction [19] of this movement strategy. Our results propose a sensory explanation for this behaviour: when perception is impaired by movement, stopping becomes a strategy to reacquire accurate sensory information. Moreover, this perception-movement coupling may apply across sensory modalities. For example, odour detection, hearing, and mechanoreception are all susceptible to motion-induced noise [12, 47-48]. And while not all animals move with intermittent motion, our simulations make further predictions about how movement may impact the perception of animals in these cases. In particular, cruise search is advantageous when perception degrades linearly with speed. Together, this supports suggestions that diverse movement strategies such as ambush and cruise search represent extremes on the same perception-driven movement-strategy continuum [16, 49], with saltatory motion an intermediate mode. Comparative work across systems, taxa and sensory modalities investigating how different perceptual constraints shape movement strategies would therefore be worthwhile.

While our study provides strong support for perceptual constraints driving saltatory motion, our experimental prey were visual stimuli that did not respond to the predator. In natural settings, prey may respond to predators, introducing feedback loops that could further shape movement strategies [50]. Moreover, perception may be impacted not only by self-induced motion, but by environmentally induced noise [29], which could further impact the movement strategies animals adopt. While the sole goal of our model was to detect prey, some movement strategies may also be more conspicuous to predators [51] or may impose different energetic costs [44]. Finally, our model used randomly distributed prey, but prey distribution may also impact the locomotory strategy of the predators, especially in case of patchy prey distributions [52]. Future work could explore how animals resolve potentially competing objectives in their movement decisions in relation to perceptual goals.

Our study provides evidence that self-induced perceptual degradation as a result of motion can be a driver of movement strategy in animals. Sticklebacks employ saltatory motion that aligns closely with the empirically derived strategy that maximizes prey detection under perceptual constraints. These results support a broader view of animal movement as not only an explorative and energetic process but also as a perceptual adaptation. Understanding this interplay is necessary to make predictions about how animals move through their environment.

