## Supporting information for "Perceptual constraints of motion shape animal movement strategies"

Supporting Information Text

**Section 1. Prey detection experiment**

1.1 **Animals**

Three-spined sticklebacks (*Gasterosteus aculeatus*) were collected using hand-nets from the River Cary, Somerset, UK in 2017. Fish were housed in 90 L tanks with continual aeration and flow-through filtered freshwater. Housing tanks contained artificial plants and plastic cylinders to provide shelter and enrichment. Fish were fed six days per week on chironomid larvae and were housed for at least one month prior to experimentation. Fish were held under a 11:13 hr light:dark cycle with temperature maintained between 13.5 - 15 °C. On the days of testing, fish were only fed after they had been tested. All procedures were approved by the University of Bristol Animal Welfare and Ethical Review Body (UIN UB/16/047).

**1.2 Perception model comparison approach**

To model the likelihood that fish would detect prey as a function of where the prey appeared in their visual field and how fish were moving, we compared binomial Generalized Linear Models (GLM) using Akaike's Information Criterion (AIC, see main text for more details). Based on the work of Hein et al. [2], we also computed an importance metric, $I_{F}$, for each term over all the different models tested using the Equation (*S1*):

$$\begin{aligned} I_{F}=\sum_{i} \boldsymbol{1}_{i,F}e^{\left( \mathrm{AIC}_{min}-\mathrm{AIC}_{i} \right)} \#\left( S1 \right) \end{aligned}$$

With $\boldsymbol{1}_{i,F}$ an indicator function returning a value of 1 if the model $i$ contains the term $F$ and 0 otherwise. $\mathrm{AIC}_{min}$ represents the minimum AIC value among all fitted models, while $\mathrm{AIC}_{i}$ the AIC value of the model $i$. We first tested which combination of angles terms ($\alpha$ and $\varphi$ included with their absolute value or with a second order polynomial factor) returned higher $I_{F}$ values (see Fig. S2A). Using the same method, we then calculated the $I_{F}$ of all main variables of the models including the selected angles (absolute value for $\alpha$ and second order polynomial for $\varphi$, see Fig. S2B). Finally, we also calculated the $I_{F}$of the interaction terms (see Fig. S2C).

The model with the smallest AIC included as fixed factors: the speed of the fish, the distance of the fish to the prey, the absolute azimuth angle ($\alpha$) between the fish’s heading and the prey, and a second-order polynomial of the elevation angle ($\varphi$) between the fish’s heading and prey. It also included interactions between distance to prey and azimuth angle ($\alpha)$, between speed and azimuth angle ($\alpha$), and between azimuth angle ($\alpha$) and the second order polynomial of the elevation angle ($\varphi$). The $I_{F}$ analysis validated this chosen model, indicating that the terms of this model were the most important to predict fish reactions, while other terms were less predictive and excluded (Fig. S2). See supplementary Dataset S1 for more information.

This model can be described mathematically with the equation *(S2)*.

|  | $I+{(k}_{s}*s)+{(k}_{d}*d)+{(k}_{\alpha}*\alpha)+(k_{\varphi}*\varphi)+(k_{\varphi^{2}}*\varphi^{2})+(d*\alpha*k_{d\alpha})+{(k}_{s\alpha}*s*\alpha)+(k_{\alpha\varphi}*\alpha*\varphi)+{(k}_{\alpha\varphi^{2}}*\alpha*\varphi^{2}),$ | *(S2)* |
| --- | --- | --- |

With *s, d,* $\alpha$*, and* $\varphi$ being respectively the standardized values of the moving speed, the distance to the prey, the absolute azimuth angle, and the elevation angle. *I* is a constant representing the intercept (value: -3.51309). Coefficients $k_{i}$ are associated with main-effect terms of variable *I*; $k_{ij}$ with interaction terms between variables *i* and *j*; $k_{\varphi2}$ and $k_{\alpha\varphi2}$ with higher-order polynomial terms involving the elevation angle ($\varphi$).

The coefficients’ values, rounded at the third decimal, were: $k_{s}=-0.819$, $k_{d}=-1.047$, $k_{\alpha}=-1.831$, $k_{\varphi}=-0.299$, $k_{\varphi^{2}}=-0.237$, $k_{da}=-0.439$, $k_{sa}=0.038$, $k_{\alpha\varphi}=-0.376$, $k_{\alpha\varphi^{2}}=0.202$.

**Section 2. Exploration assay**

**2.1 Animals**

Three-spined sticklebacks were collected using hand-nets from streams connected to the River Cam, Cambridge, UK, in December 2020. After collection, fish were housed in aquaria (120 x 45 x 50 cm; L x W x H). Ambient temperature (12°C), light (12:12hr (L:D) and water circulation (1000 L/h) were controlled throughout housing. Aquaria were enriched with gravel substrate, artificial and real plants, and received three frozen cubes of bloodworms (12.5 g, Superfish ©) daily. These fish were maintained in the laboratory for three months prior to experimentation. During the experiment, a cohort of fish (n = 12) were transferred to another aquaria (100 x 100 x 15 cm; length x width x height) and individuals were isolated from each other using mesh containers (15 x 20 x 10 cm; length x width x height) and numbered uniquely to allow for individual identification. The aquaria conditions were identical to the housing aquaria, and each mesh container contained an artificial plant for enrichment. Each individual fish was fed five bloodworms a day. All procedures were approved by the Institutional Animal Care and Use Committee (IACUC Protocol number Z0080/20).

**Section 3. Perceptual model**

**3.1 Degradation of perception with increasing speed**

To characterize the general shape of the degradation in perception with increasing speed, we superimposed a 2 m radius sphere centred on the fish and calculated the probability of detection per frame for all possible locations within the sphere, using a 2 cm grid system to define the coordinates of each location. We chose a radius of 2 m because the perception field drawn by the GLM model showed that prey from further away were unlikely to be detected (see Fig. S3). To generate an overall probability of prey detection, the probability at each location was averaged across the grid. This process was repeated across various speeds, ranging from 0 to 450 mm/s with increments of 10 mm/s. To obtain the perception degradation ratio (as seen in Fig. S4), we divided each probability value (at each speed) by the probability value obtained with a speed of 0 mm/s.

**3.2 Agent parameters and environment**

We built a three-dimensional spatially explicit agent-based model in Python (version 3.9) to simulate the differential movement patterns of fish when searching for prey along a straight trajectory line. Each simulation involved an individual agent moving along this line for three minutes (timestep $\Delta_{t}$= 1/23.98 sec). At each timestep, the agent could either stop or be in motion and its perceptual field is updated according to our perception model and perception degradation ratio. We chose a timestep of 1/23.98 sec to align with the frame rate of 23.98 fps of the camera used in the exploration assay. Prey positions were randomly assigned along the trajectory line, and the total number of prey was calculated to achieve a density of 0.5 prey per m³, facilitating a direct comparison between model predictions and experimental data. Before each simulation, we specified i) the maximum speed of the agent while moving (S), ii) the average duration of an agent’s stopping phase ($\Delta_{\mathrm{stop}}$), iii) the average duration of an agent’s motion bout ($\Delta_{\mathrm{move}}$), and iv) the effect of speed on the agent’s perceptual field.

**3.3 Agent movement**

We defined a motion model that captured the features of the fish’s movements from the empirical data extracted from the exploration assay (see main text). At each timestep, if an agent stopped, it had a probability $P_{stop \to move}= \frac{1}{\Delta_{\mathrm{stop}}/ \Delta_{t}}$ of transitioning to an acceleration phase, ensuring that the average duration of stopping phases was equal to $\Delta_{\mathrm{stop}}$. The duration of the acceleration phase varied according to the speed the fish adopted during movement (see below). After the acceleration phase, the agent entered a movement phase, during which it had a probability$P_{move \to stop}=\frac{1}{\Delta_{\mathrm{move}}/ \Delta_{t}}$of switching to a deceleration state. Like the acceleration phase, the duration of the deceleration phase varied according to the speed the agent adopted while moving and ended with the agent entering a stopping phase. If the agent was in a movement phase during a time step, its position was modified such that it moved forward according to its speed.

**3.4 Acceleration and deceleration patterns**

The analyses of the exploration assays suggested that the fish needed a minimum time to accelerate to their moving speed, and a time for deceleration after this movement phase ended. We chose to model these two phases based on the experimental data. To achieve this, we extracted all the events of acceleration and deceleration phases from the experimental data (see main article and Fig. S8) and normalized these values so that each acceleration and deceleration event ranged from 0 to 1 in both the amplitude and time axes. We then randomly selected a subset of 10,000 acceleration and 10,000 deceleration events (out of 88,001 and 87,271, respectively), excluding any of a duration that was smaller than 3 frames (125 milliseconds). We then fitted these values with sigmoidal functions of the equation: $\frac{\theta}{\left( 1+\beta*\exp\left( \gamma*t \right) \right)}$ with *t* representing normalized time using the R integrated *optim* function. We found the parameters for the shape of the acceleration pattern: θ = 1.067, β = 33.554, and γ = -5.853; and for the deceleration pattern: θ = 1.010, β = 0.061, and γ = 5.707 (see Fig. S10 A-B). The durations of these phases were positively correlated with the maximum speed reached during the associated movement phase. To determine the duration of these acceleration and decelerations phases, we again fitted experimental data with the *optim* function. This time, we fitted the data with an asymptotic function $(a-\left( a-b \right)*\exp\left( -c*S \right)$ with *S* representing the maximum movement speed of an associated movement phase. We obtained *a* = 0.416, *b* = 0.091, *c* = 0.068 for accelerations, and *a* = 0.658, *b* = 0.022, *c* = 0.066 for decelerations (see Fig. S10 C-D). From these functions, we were able to calculate, for the chosen movement speed, the duration and shape of the acceleration and deceleration phases, and these were implemented in the agent-based model.

**3.5 Probability of prey detection.** At each timestep, we calculated the probability that a fish would detect a prey. If the distance to the prey exceeded 2 m, then this probability was set to 0. If the distance to the prey was within 2 m, we calculated its probability of detection for one timestep ($\Delta_{t}$), $P_{\mathrm{detect}\Delta_{t}}$, using the following equation *(S3)*:

$\begin{aligned} P_{\mathrm{detect}\Delta_{t}}= 1- {exp}^{\frac{\ln\left( 1-P_{\mathrm{detect}\Delta_{T}} \right)*\Delta_{t}}{\Delta_{T}}} ,\#\left( S3 \right) \end{aligned}$

where $P_{\mathrm{detect}\Delta_{T}}$ is the probability of detection predicted from experimental data over a given Δ_T_ duration (Δ_T_ = 0.88 seconds, the duration of the prey presentation). We used a GLM model to generate the $P_{\mathrm{detect}\Delta_{T}}$ value, which also accounted for the movement speed of the fish, the azimuth and elevation angles of the fish’s heading to the prey (*SI Appendix*, Section 1.2 and Equation S2, and Fig. 1D), and the distance between the fish and the prey. We then used these values to determine whether each prey was detected by the fish. Several prey could be detected during the same timestep. When a given prey was deemed to have been detected, they were removed from the simulation. For simplicity, the presence or absence of prey did not influence the movement parameters of the fish.

We also aimed to see how the optimal movement strategies changed according to how movement speed impacted the loss of perception. Anderson (5) hypothesized that a saltatory search was the optimal strategy if an organism suffers from a convex-shaped decrease in perception with speed, but not for a linear decrease in perception (see Fig. S4). To evaluate whether this applied to the present study, we conducted a second set of simulations in which the probability of prey detection was adjusted using a linear-to-convex ratio. This approach ensured that the overall probability of detection followed a linear decline, while preserving the relative differences in perceptual field shape. For example, as speed increases, this method maintains the reduction in peripheral vision relative to frontal vision, while enforcing an overall linear decrease in perception. To calculate the probability for a given prey to be detected under linear perception model, we used the following equation *(S4)*:

$$\begin{aligned} P_{\mathrm{detect}_{linear\Delta t}}= P_{\mathrm{detect}\Delta_{t}}* \frac{P_{\mathrm{linearS}}}{P_{\mathrm{convexS}}} = P_{\mathrm{detect}\Delta_{t}}* \frac{1 -0.002002428 * S}{{0.9948719}^{S}}. \#\left( S4 \right) \end{aligned}$$

Here, S represents the movement speed, while $P_{\mathrm{linearS}}$ and $P_{\mathrm{convexS}}$ represent the estimated overall probability of prey detection obtained for a linear decrease in perception and for a convex decrease in perception (as in Fig. S4), respectively. The parameters for the exponential decrease part of this equation were found using the *optim* function in R to fit the data obtained, and as described above. In this case, we used a linear decrease starting at P = 1 and reaching at 450mm/s the same detection probability as the one of the convex model prediction at 450 mm/s. We also ensured that other kinds of linear decrease would have given similar results (see supplementary Fig. S9).

Lastly, we also implemented a model to reflect a ‘blindness’ for fish when moving. To achieve this, the probability of prey detection when agent was in a stopping phase reflected the probability predicted by our GLM model but, when moving, this probability was then set to 0 (see Fig. S4).

Figures

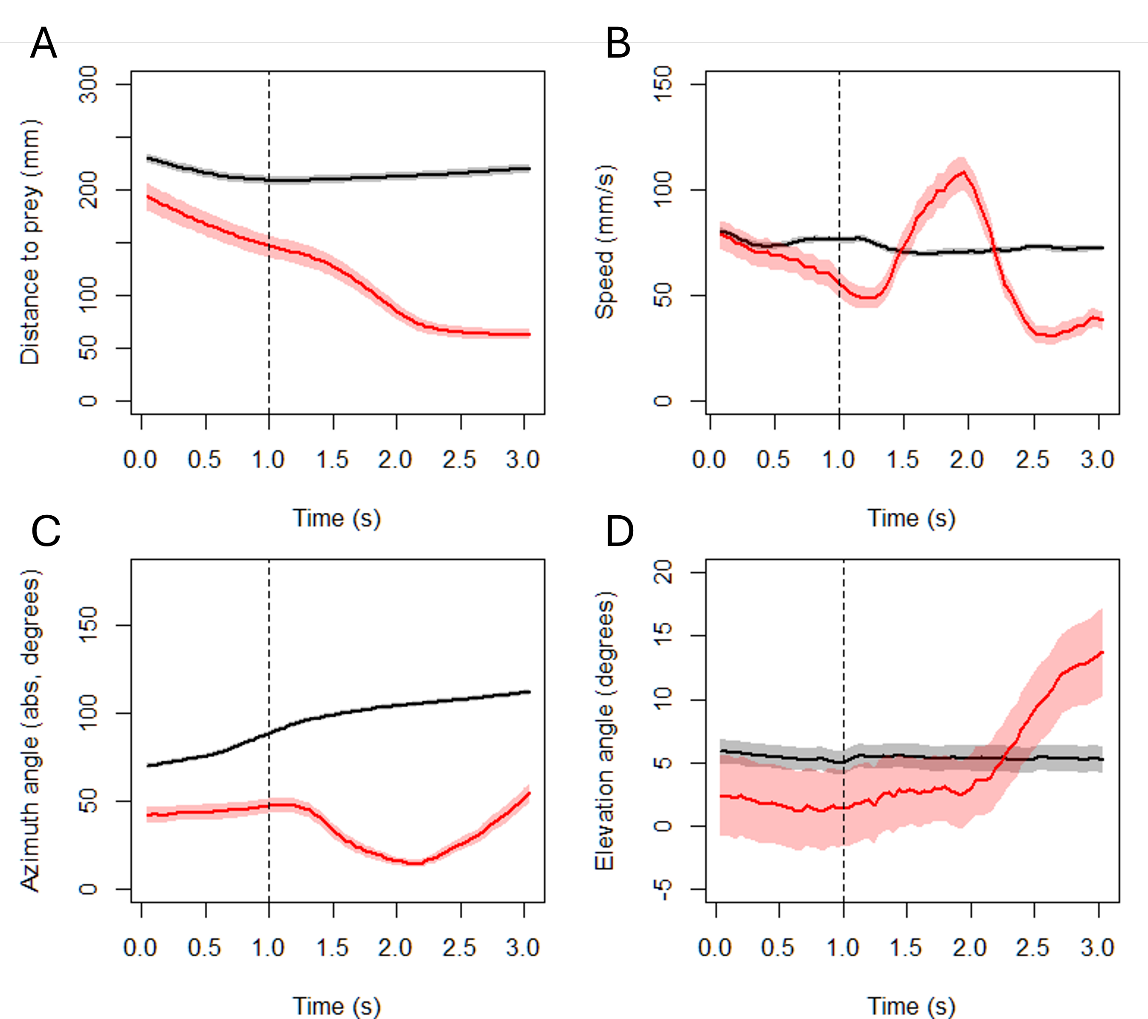

Fig. S1. Quantification of the differences in behaviour between fish who reacted to the prey presentation (red) and fish who did not react to the prey presentation (black). A) Changes in distance between fish and prey over time, B) Changes in fish self-induced motion speed, C) changes in absolute value of Azimuth angle *α*, D) changes in elevation angle *φ* over time. The dotted vertical black line indicates the moment the prey appears. The difference between reactive and non-reactive fish before 1 second illustrates how the difference in behaviour or location relative to prey can impact the probability of reaction to prey. The strong changes in behaviour occurring in the 2 seconds following prey apparition confirm that fish identified as “reacting to the prey” respond differently to the prey compared to non-reactive fish. Shaded areas indicate 95% CI.

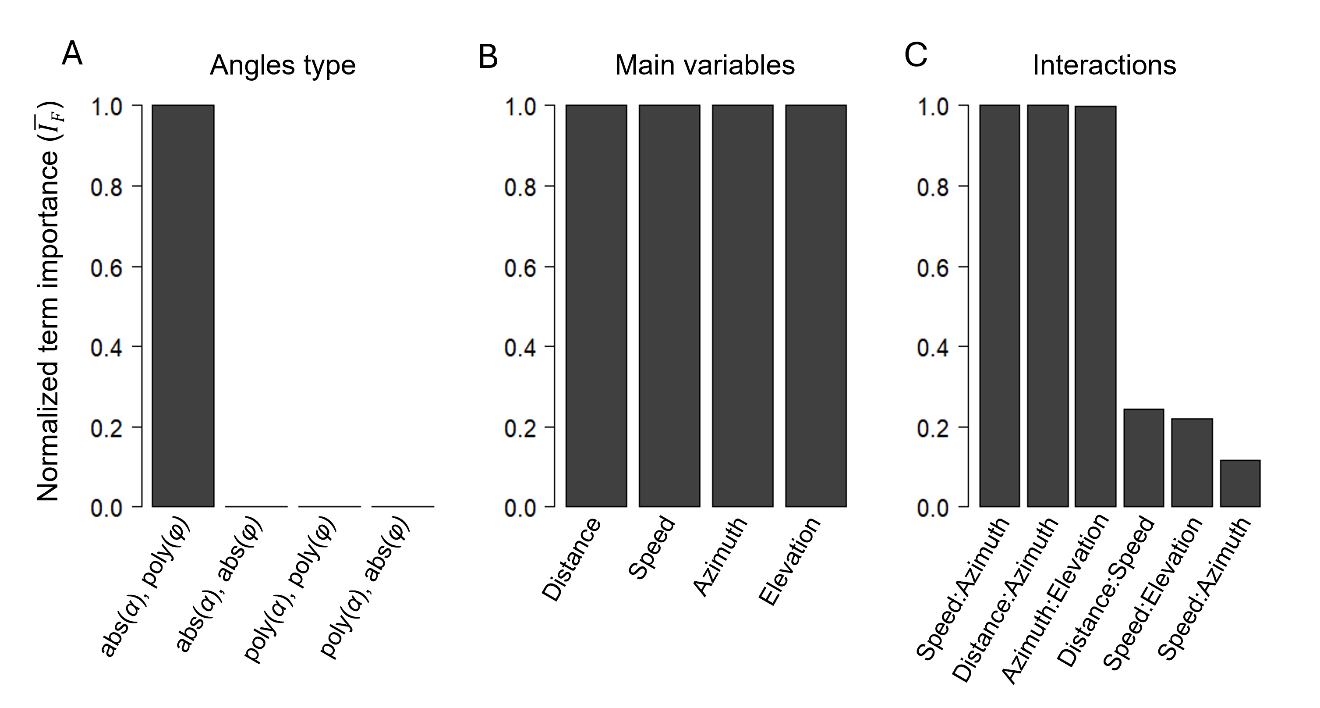

Fig. S2. Importance of the terms included in the different GLMs used to model the fish’s visual perception. The Importance Factor $\boldsymbol{I}_{\boldsymbol{F}}$ was calculated according to the Equation S1 and scaled so that the maximum value within each panel equals 1. A) Comparison of $\boldsymbol{I}_{\boldsymbol{F}}$ values for models including different type of angular variables (azimuth $\boldsymbol{\alpha}$ and elevation $\boldsymbol{\varphi}$). The notation abs(angle) refers to the absolute value of the given variable, while poly(angle) to the usage of a second-order polynomial term. B) Importance of the four main variables when using the absolute value of azimuth and a second order polynomial for elevation. C) As in panel B, this panel shows the importance of the different interaction terms. These importance values validate the final selected model, which included the four main variables (absolute azimuth values and elevation as second order polynomial variable), as well as the three most relevant interactions: Speed x Azimuth, Distance x Azimuth, and Azimuth x Elevation.

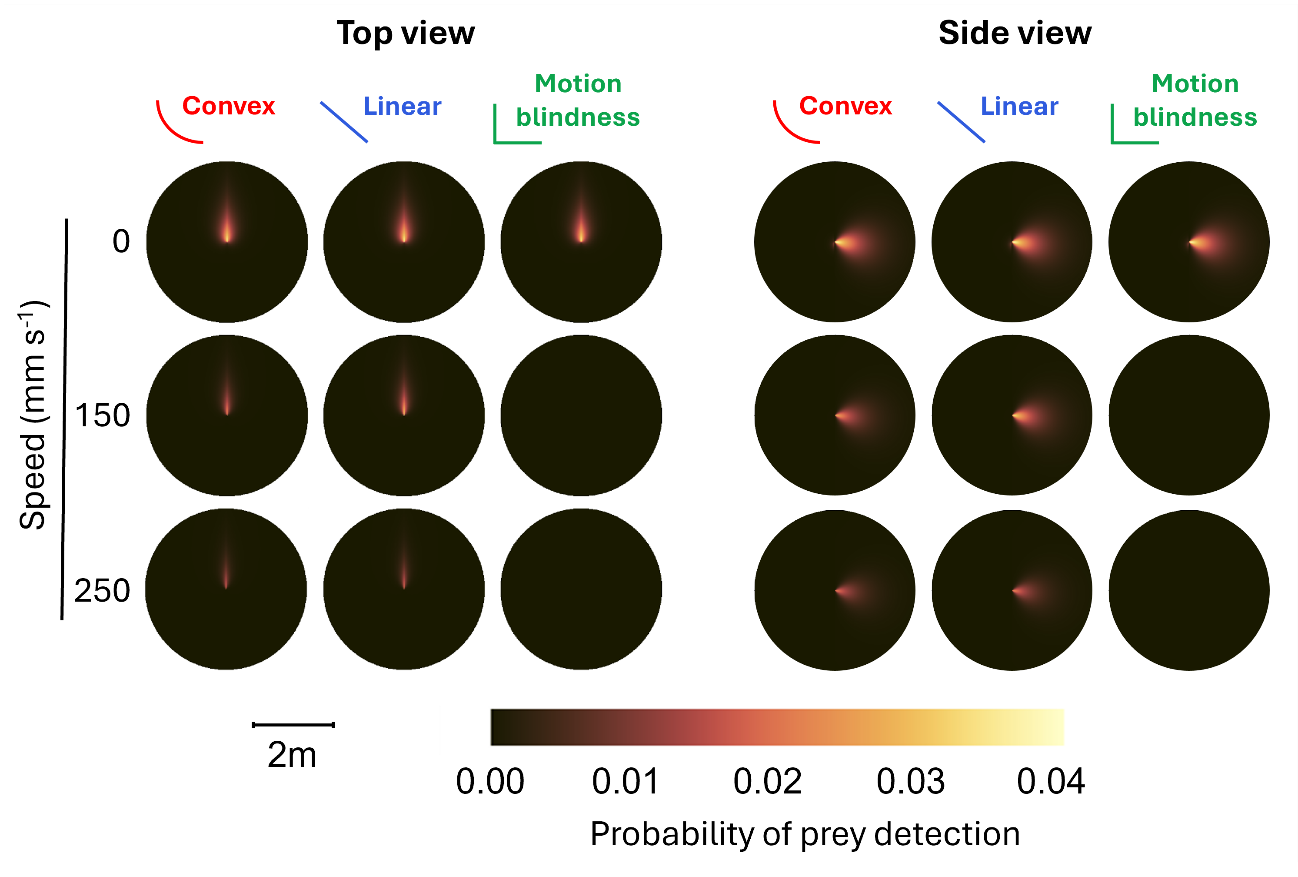
Fig. S3. Illustration of the perception map across the horizontal plane (top view) and across the vertical plane (side view) for three different models of perception decrease when moving: convex decrease, linear decrease and motion blindness (see Fig. 2A). In all three models, the perception map is similar for a speed of 0. For the convex model, the overall perception rapidly decreases with an increase in speed, whereas, for the linear model, perception decreases more gradually as speed increases. For the motion blindness model, the agent does not perceive anything when it moves, irrespective of speed.

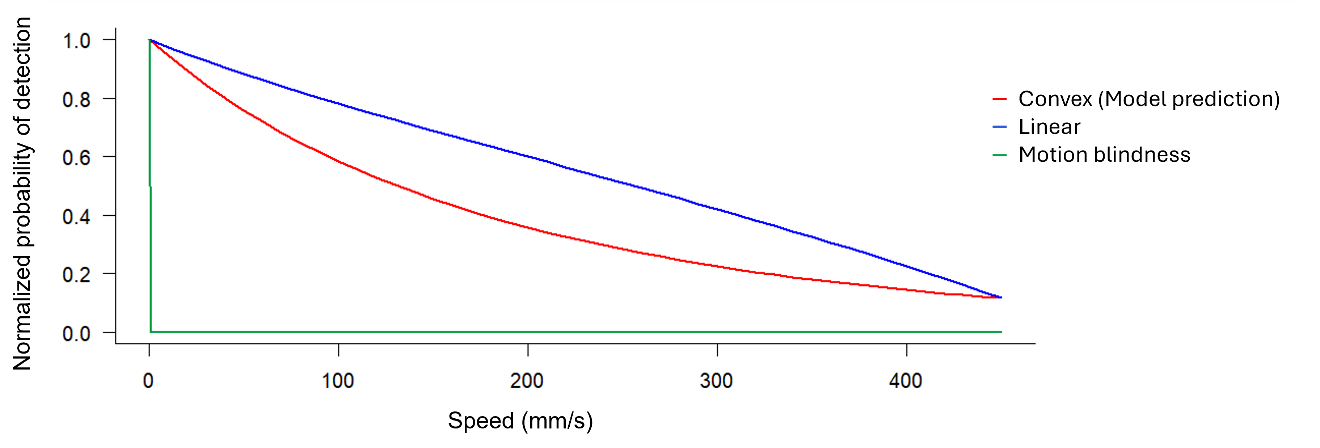

Fig. S4. Predicted overall probability of detecting a prey within a 2 m radius sphere as a function of the speed of the fish. The red line represents values predicted by the GLM model (see *SI Appendix*, section S1.4 and Equation S2). Two different models are also presented, as used in the simulations. In blue is the function used to simulate a linear decrease in the probability of detection as a function of speed. In green is the function used to mimic the effect of motion blindness.

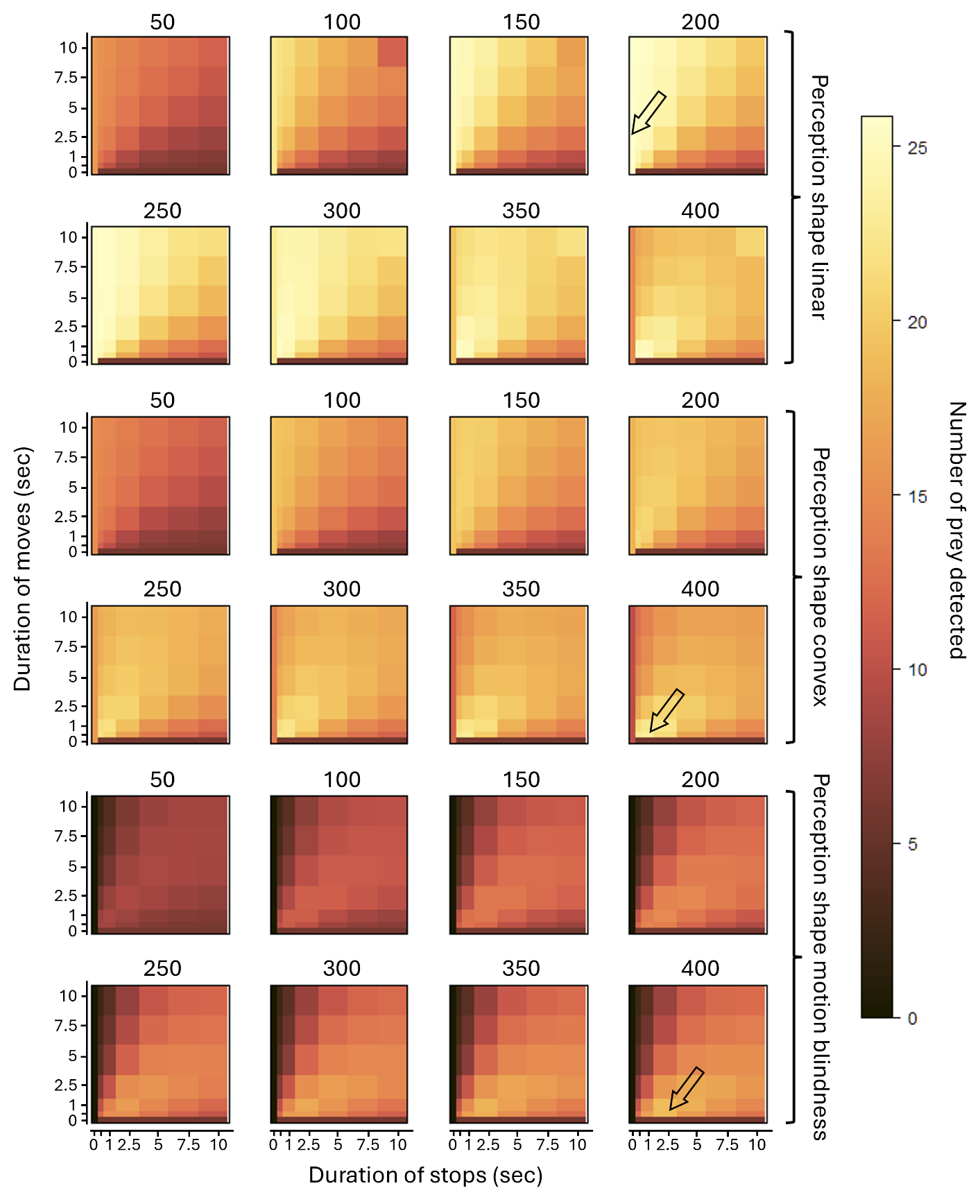

Fig. S5. Heatmaps showing how the average number of prey detected during simulations (3 mins each; N = 5,000) changed as a function of movement duration, stop duration, movement speed, and the model of perceptual decline (either linear, convex, or motion blindness). The value above each heatmap represent the maximum moving speed of the agent (mm/s). The same 5,000 prey presentations were used for each grid cell (defined by the axis ticks) to limit uncontrolled variation between conditions. The black arrows have been added because the variation in colour may be difficult to distinguish visually: the arrows indicate the position of the simulation with the maximum number of detected prey for the associated perceptual function.

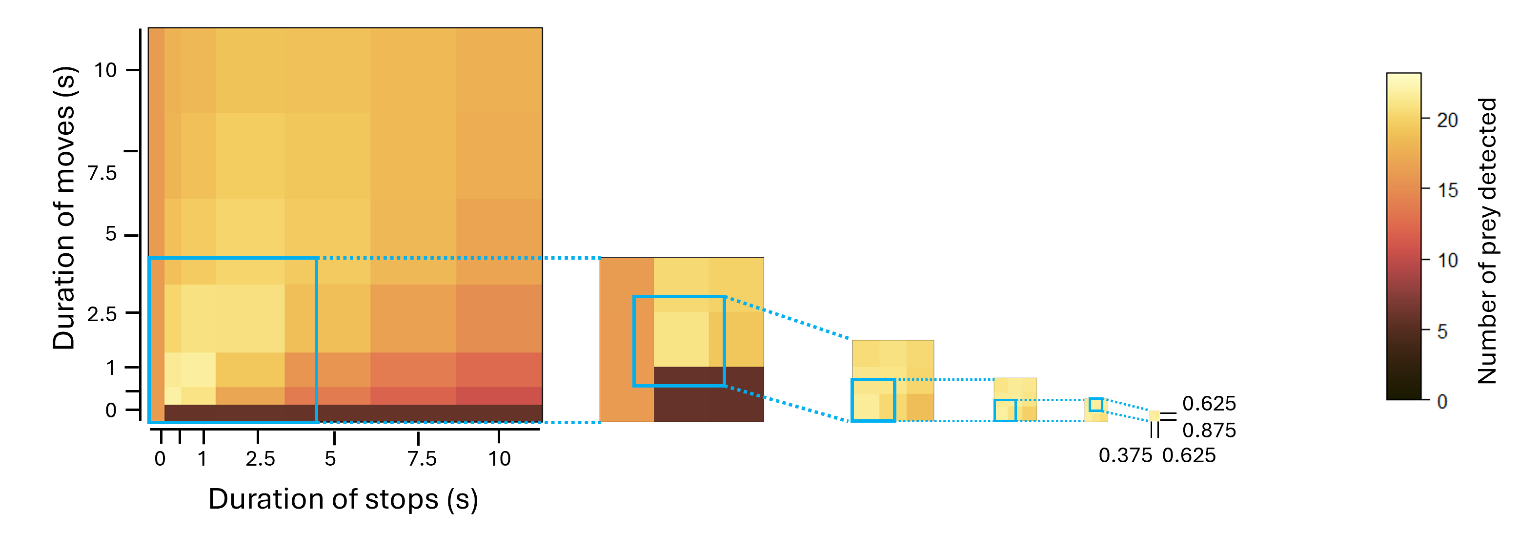

Fig. S6. Illustration of the iterative grid refinement process to find the optimal movement strategy in simulations. The heatmaps show how the average number of prey detected during simulations (3 mins) varied as a function of the duration of movements and stops. The same 10,000 prey presentations were used for each grid cell to limit variation between conditions. The original heatmap (far left, Fig. S4) was generated using 5,000 replicates per parameter combination, with stop and movement durations of 0 s, 0.5 s, 1 s, 2.5 s, 5 s, 7.5 s, and 10 s. Subsequent grid refinements used N = 10,000 replicates. The first refinement tested nine parameter combinations centred on the previously best-performing cell at a 2 s resolution (e.g., 0 s, 2 s, 4 s). Each following iteration halved the grid resolution (1 s, 0.5 s, 0.25 s, etc.), with the final iteration reaching 0.125 s.

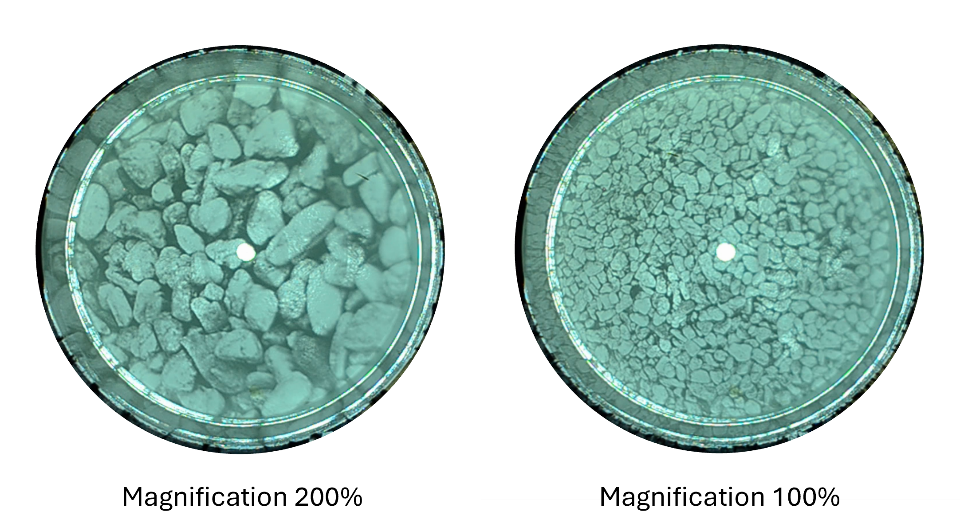

Fig. S7. Example images of the two backgrounds used in the exploration assay experiment.

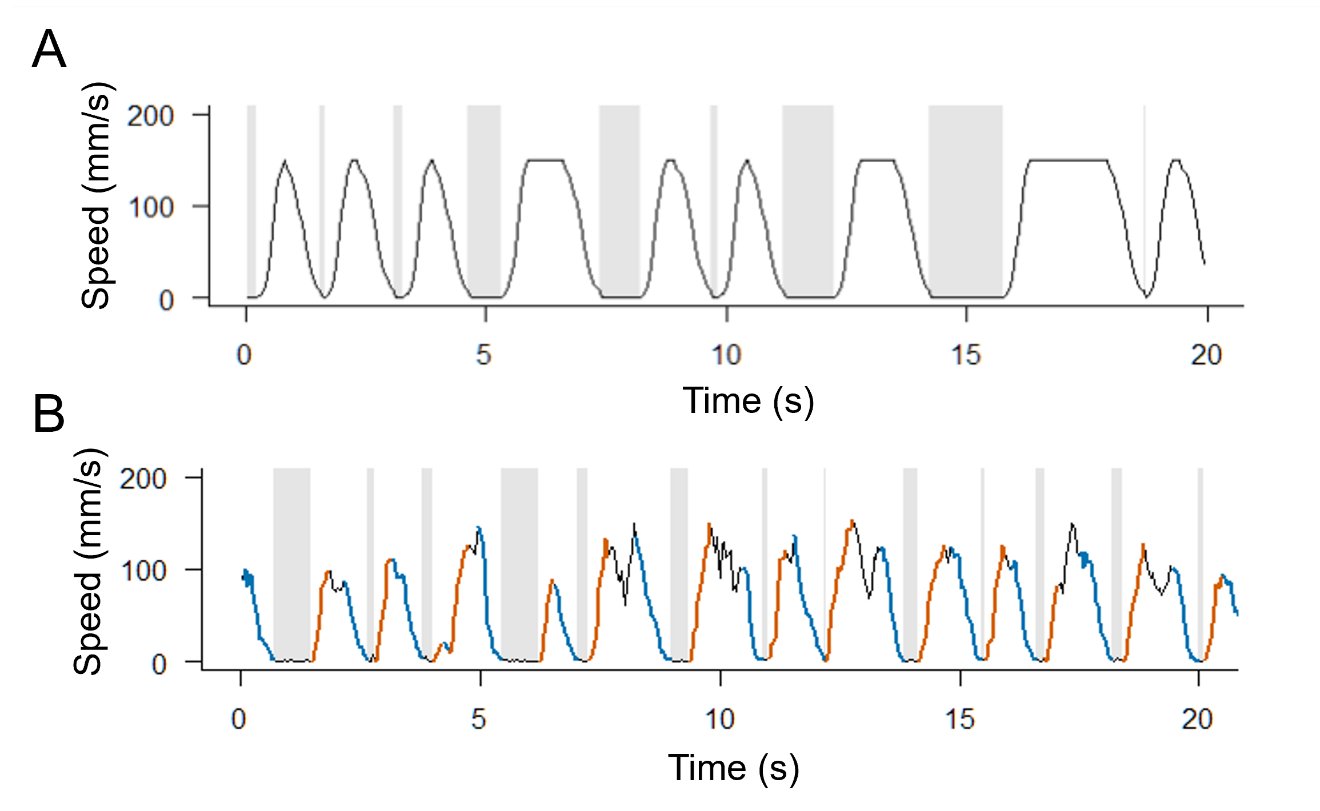

Fig. S8. Example of a simulated speed profile generated by the model (A) and of a stickleback exploring a circular arena (B). The simulation was conducted with an average stop duration of 1 s and an average movement duration of 0.5 s, with a maximum movement speed of 150 mm/s. Shaded areas represent stops, while non-shaded areas indicate movement. These parameters were adjustable within the model, allowing for customized simulations. For the speed profile from empirical data, the shaded areas indicate the parts identified as stops by our algorithm (see *SI Appendix*, section S2.4), while non-shaded regions are movements. Orange or blue sections (plot on the right) indicate phases identified as accelerations or decelerations, respectively.

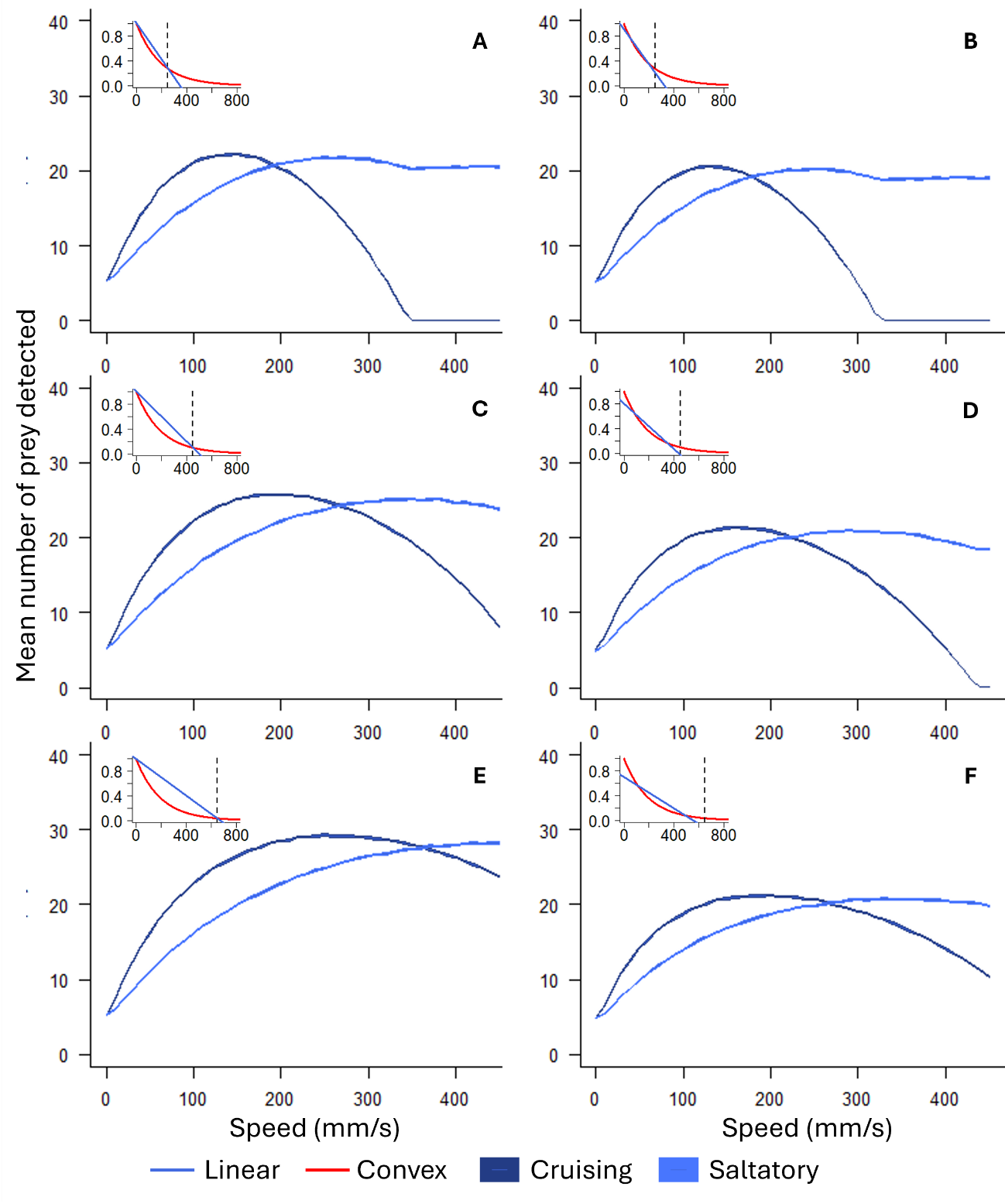

Fig. S9. Average number of prey detected in simulations (3 mins each; the maximum speed was varied in steps of 10 mm/s, and N = 5000 repetitions were performed at each step) when moving at different maximum speeds and in two different ways: cruising (constantly moving; dark blue line) and saltatory (1s average stop duration and 1s average movement duration; light blue line) for various linear perceptual decrease models (inset plots). Line width represents 95% confidence intervals. There are various ways of calculating the linear decrease in perception as presented in the sub-panels. In those sub-panels, the red line represents the convex function obtained from our empirically defined model and is represented as an indicator. We illustrate that the method used to calculate linear perceptual decrease does not alter the general result that, under linear decrease, cruising outperforms saltatory motion when comparing the best performance among all speeds. Even though saltatory strategy may reach very similar values of prey detection at high maximum moving speeds, they remain statistically lower than the ones obtained with cruising strategy (for all cases: Welch modified two-sample t-test: P < 0.001, 9985 < df < 9997.9, 4.02 < t < 9.9). For example, panels A, C, and E illustrate use of a linear decrease function that has an intercept at 1.0 and which calculates the slope so that both the convex perception function and linear perception function cross at a given speed (indicated by the vertical dashed lines in sub-panels). Alternatively, the other panels (B, D and F) illustrate a linear decrease function that has an intercept and slope that are calculated such that the area under each curve is similar for both convex and linear decrease for speed (below a given speed value; indicated by the vertical dashed lines in corresponding sub-panels). The linear decrease function used in the current study is represented in panel C.

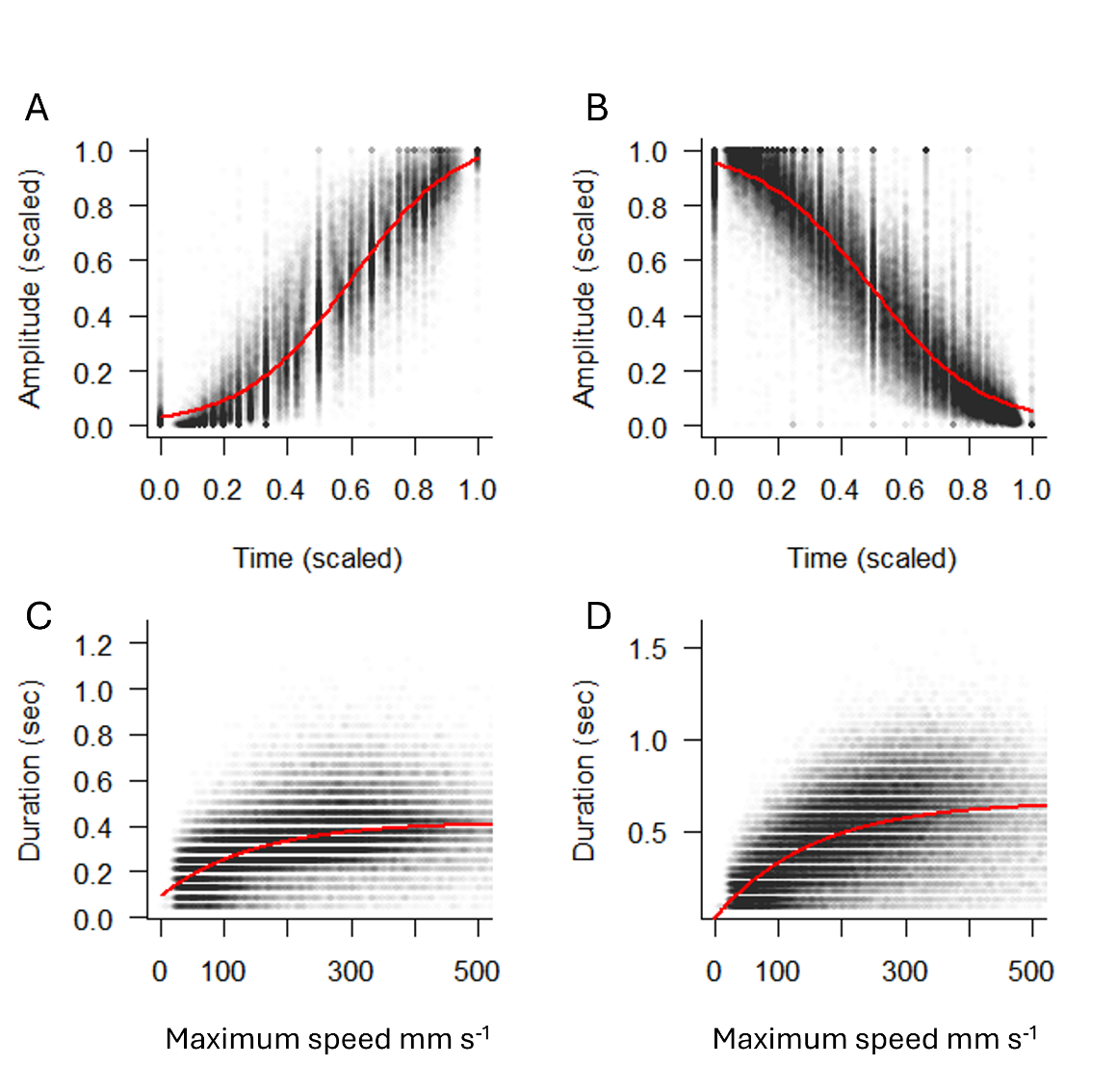

Fig. S10. A-B) Acceleration (A) and deceleration (B) normalized data with the fitted function used in the simulations. C-D) Duration of acceleration (C) and deceleration (D) phases plotted against the maximum speed reached during the corresponding movement phases. Red lines indicate functions used to fit these data and to mimic empirical acceleration and deceleration patterns in the simulations.

Tables

Table S1. Parameter estimates (± standard error) and significance levels (p-values) from the best-fitting binomial generalized linear model (GLM) predicting the likelihood of prey detection in the prey detection experiment. The model includes fish movement speed, fish-prey distance, azimuth ($\boldsymbol{\alpha}$), and elevation ($\boldsymbol{\varphi}$), and their interaction terms. Significance codes: *** p < 0.001; ** p < 0.01; values without stars indicate non-significant effects.

|  | **Estimate ± s.e.** | | | z value | P |  |
| --- | --- | --- | --- | --- | --- | --- |
| Fish movement speed | -0.819 | ± | 0.125 | -6.565 | <0.001 | *** |
| Fish-prey distance | -1.047 | ± | 0.146 | -7.157 | <0.001 | *** |
| Azimuth $\alpha$ | -1.831 | ± | 0.154 | -11.876 | <0.001 | *** |
| Elevation $\varphi$ (1st order) | -0.299 | ± | 0.089 | -3.383 | <0.002 | *** |
| Elevation $\varphi$ (2nd order) | -0.237 | ± | 0.065 | -3.658 | <0.001 | *** |
| Fish-prey distance * azimuth $\alpha$ | -0.439 | ± | 0.138 | -3.183 | 0.001 | ** |
| Movement speed * azimuth $\alpha$ | -0.376 | ± | 0.113 | -3.330 | <0.001 | *** |
| Azimuth $\alpha$ * elevation $\varphi$ (1st order) | 0.038 | ± | 0.087 | 0.440 | 0.660 |  |
| Azimuth $\alpha$ * elevation $\varphi$ (2nd order) | 0.202 | ± | 0.063 | 3.187 | 0.001 | ** |

**Movie S1 (separate file).** Representation of the regions around a fish where the probability to detect a prey within the shaded volume is 0.15 or higher. The shape and volume of this field varies according to the fish’s speed. The schematic of the fish indicates a fish’s position and orientation, to scale.

**Dataset** **S1** **(separate file).**

**Sheet “Prey detection expe. data”. Empirical data used to model the perception field of the fish. The table contains the spatial coordinates and movement speed of the fish relative to prey at the moment the prey appears.**

**Sheet “Perception models”. List of all the GLMs tested to model the perception field of the fish. All possible model combinations are included with corresponding AIC values.**

**Sheet “Exploration assay expe. data”. Empirical data characterizing fish motion behavior during the exploration assay.**

**Sheet “Simulations for Fig. 4B”. Simulation data used to examine the effects of speed on the shape of perceptual decrease.**

**Sheet “Simulations for Figs 4C, S2”. Simulation data used to identify which motion strategy maximized the number of detected prey according to the shape of perceptual decrease and speed limitations.**

**Sheet “Simulations for Fig. S3”. Simulation data used to compare the effect of different forms of linear perceptual decrease on prey detection probability according to fish movement speed.**

**More detailed description of each sheet is provided within the Excel file itself.**
